# Quantitative assessment of eye phenotypes for functional genetic studies using *Drosophila melanogaster*

**DOI:** 10.1101/036368

**Authors:** Janani Iyer, Qingyu Wang, Thanh Le, Lucilla Pizzo, Sebastian Grönke, Surendra S. Ambegaokar, Yuzuru Imai, Ashutosh Srivastava, Beatriz Llamusí Troisí, Graeme Mardon, Ruben Artero, George R. Jackson, Adrian M. Isaacs, Linda Partridge, Bingwei Lu, Justin P. Kumar, Santhosh Girirajan

**Affiliations:** Department of Biochemistry and Molecular Biology, The Pennsylvania State University, University Park, PA 16802; Bioinformatics and Genomics Program, Huck Institute of Life Sciences, The Pennsylvania State University, University Park, PA 16802; Max-Planck Institute for Biology of Aging, Cologne, Germany; Department of Botany and Microbiology, Ohio Wesleyan University, Delaware, OH 43015; Department of Research for Parkinson's Disease, Juntendo University Graduate School of Medicine, Tokyo, Japan; Translational Genetics Group, Department of Genetics, University of Valencia, Burjassot, Spain; Incliva Health Research Institute, Avda. Menéndez Pelayo, 4 accesorio, 46010 Valencia. Spain.; Departments of Pathology and Immunology and Molecular and Human Genetics, Baylor College of Medicine, Houston, TX 77030; Department of Neurology, Baylor College of Medicine, Houston, TX 77030; Department of Neurodegenerative Disease, UCL Institute of Neurology, London, WC1N 3BG, UK; Department of Pathology, Stanford University, Palo Alto, CA; Department of Biology, Indiana University, Bloomington, IA 47405; Department of Anthropology, The Pennsylvania State University, University Park, PA 16802

**Keywords:** Ommatidia, *Drosophila melanogaster*, neurodevelopmental disorders, modifier screens, rough eye, human disease models

## Abstract

About two-thirds of the vital genes in the *Drosophila* genome are involved in eye development, making the fly eye an excellent genetic system to study cellular function and development, neurodevelopment/degeneration, and complex diseases such as cancer and diabetes. We developed a novel computational method, implemented as Flynotyper software (http://flynotyper.sourceforge.net), to quantitatively assess the morphological defects in the *Drosophila* eye resulting from genetic alterations affecting basic cellular and developmental processes. Flynotyper utilizes a series of image processing operations to automatically detect the fly eye and the individual ommatidium, and calculates a phenotypic score as a measure of the disorderliness of ommatidial arrangement in the fly eye. As a proof of principle, we tested our method by analyzing the defects due to eye-specific knockdown of *Drosophila* orthologs of 12 neurodevelopmental genes to accurately document differential sensitivities of these genes to dosage alteration. We also evaluated eye images from six independent studies assessing the effect of overexpression of repeats, candidates from peptide library screens, and modifiers of neurotoxicity and developmental processes on eye morphology, and show strong concordance with the original assessment. We further demonstrate the utility of this method by analyzing 16 modifiers of *sine oculis* obtained from two genome-wide deficiency screens of *Drosophila* and accurately quantifying the effect of its enhancers and suppressors during eye development. Our method will complement existing assays for eye phenotypes and increase the accuracy of studies that use fly eyes for functional evaluation of genes and genetic interactions.

## Article Summary

We present a quantitative tool, Flynotyper, for assessment of eye phenotypes for functional studies in *Drosophila melanogaster*. Using proof-of-principle experiments we quantify dosage sensitivity of 12 neurodevelopmental genes, accurately validate our method across different imaging platforms and genotypes from six independent studies, and demonstrate the utility of our tool by analyzing novel modifiers of *sine oculis (so)* obtained from two genome-wide deficiency screens and classifying them into enhancers and suppressors based on their effect on so-associated eye phenotypes. Our method will complement existing assays for eye phenotypes and increase the accuracy of studies that use fly eyes for functional evaluation.

## Introduction

Current strategies for functional analysis of genes in various biological processes using animal models have been limited due to the lack of highly sensitive and quantitative assays. *Drosophila melanogaster* remains a powerful model for genetic studies with about 75% of human disease genes having orthologs in flies (1). *Drosophila* provides a wealth of genetic, cellular, and molecular biology tools, which have been instrumental in understanding basic biological processes (2). With the availability of such tools and high conservation of human disease-associated genes, the past decade has seen the growth of *Drosophila* models to study human diseases (3). Specifically, the fly eye is an excellent experimental system for high throughput genetic screening and in dissecting molecular interactions (4). Two-thirds of the vital genes in the *Drosophila* genome have been estimated to be required for eye development (5). Although some genes are likely to be specific for eye development, other vital genes expressed in the eye are probably required for general cellular processes as well (4). Hence, phenotypic assessment of the eye can be extended to gene functions in other tissues. Since it is a dispensable organ for survival, studies using the fly eye have been used for understanding basic biological processes including cell proliferation and differentiation, neuronal connectivity, apoptosis, and tissue patterning (6).

The *Drosophila* compound eye is a simple nervous system consisting of a symmetrical organization of approximately 750 ommatidia (7). Each ommatidium contains eight photoreceptor neurons orchestrated in a trapezoid fashion and surrounded by four lens-secreting cone cells and two primary pigment cells. The ommatidia are separated from one another by a lattice of twelve accessory cells that include six secondary pigment cells, three tertiary pigment cells, and three mechanosensory bristle complexes (8). Since the structure of the eye is ordered precisely, any subtle defect that alters the geometry of a single ommatidium or disrupts the development of a single cell within the ommatidium leads to observable morphological phenotypes such as the rough eye. Other commonly observed eye phenotypes can include small or large eye, change in size of individual ommatidia, changes in bristles and loss of pigmentation. Genetic screens for modifiers of a phenotype caused by knockdown/mutation or misexpression of a gene in the developing eye have played a pivotal role in identifying novel genes interacting in the same or different biological pathways (9, 10). The majority of genetic screens utilizing the eye phenotypes take advantage of the rough eye or changes in the size of eye. The rough eye phenotype could arise due to lack of individual photoreceptor neurons or change in the number, arrangement or identity of photoreceptor neurons (11-14). *Drosophila* rough eye phenotypes have been utilized to identify genetic modifiers of genes including *ras* (6), *ksr* (15), *sina* (16), *sine oculis* (17), and humanized models of *MECP2* (10) and *ATXN3* (18). However, these studies assessing the rough eye morphology are qualitative in nature and hence open to varied interpretations. Usually, the different eye phenotypes are visually analyzed and manually rank ordered based on their severity. While severe, overt eye phenotypes are readily recognizable to the naked eye, differentiating subtle alterations can be challenging. In the modifier screens, while strong enhancers and suppressors can be identified by qualitative analysis (i.e. visual inspection) of the eye phenotypes, weak modifiers may go undetected. Currently, no image analysis techniques are available that can automatically and accurately quantify the rough eye phenotypes observed in bright field or SEM images.

Here, we present a novel computational method, which facilitates accurate analysis of *Drosophila* rough eye morphology from images obtained using bright-field microscopy or SEM. Using morphological transformation to detect the fly eye and ommatidial measurements to quantify the disorderliness, this sensitive assay can detect morphological changes in *Drosophila* eyes. We tested our automated method by analyzing the morphological defects resulting from eye-specific knockdown of *Drosophila* orthologs of 12 genes known to be associated with neurodevelopmental disorders. We also validated our method by analyzing a representative group of eye images from six independent published and unpublished studies, including a genetic screen for modifiers of *tau*-induced neurotoxicity (19), screen for interactors of *dFoxO* (20), screen for interactors of *DJ-1*, peptide library screen for *Drosophila* model of myotonic dystrophy type 1 (DM1) (21), and a genetic screen for interactors of *Egfr* (22). We also validated the roughness of the fly eyes overexpressing *C9orf72* pure repeats and RNA only (RO) repeats (23). Our quantitative analysis and classification of these modifiers using Flynotyper was concordant with the authors’ original qualitative assessment. We further used this method for assessing the genetic modifiers of *sine oculis (so)*, a key gene involved in eye formation, obtained from two genome-wide screens and accurately classifying their effect on the *so*-associated eye phenotype. Our method provides a quantitative tool for dissecting phenotypic heterogeneity frequently observed in studies of genetic mutations, gene dosage alterations, and genetic modifiers in development and disease.

## Results

### Automated detection of fly eye and calculation of phenotypic score

We have developed an automated computational method to quantify observable morphological defects in the *Drosophila* eye from images taken using bright-field microscopy or SEM. The algorithm is implemented as Flynotyper software and is available as an open-source package (http://flynotyper.sourceforge.net/) and ImageJ plugin (**Figure S1**). For each image, a series of morphological transformations were first applied to suppress the image background and identify the fly eye (24) (**Figure 1A-E**). Then each ommatidium was isolated and its center localized using image transformation and searching for local maxima (**Figure 1F-I**). Based on the hexagonal arrangement of ommatidia, we determined groups of six local vectors directed from the center of each ommatidium to the centers of six neighboring ommatidia (**Figure 2**). We quantified the disorganization of the ommatidia by applying the principle of entropy, a measure of disorderliness, in which a perfectly radial symmetry will result in zero entropy. We calculated ommatidial disorderliness indices (ODI) of a fly eye using the cumulative differences in the lengths and angles formed by these adjacent local vectors (see Methods). Phenotypic scores were then determined using the ODI and the estimated number of fused ommatidia (fusion index) (**Figure S2**). A higher phenotypic score represents increased disorderliness or altered symmetry of the ommatidial arrangement and thus increased severity of the eye phenotype.

**Figure 1.**
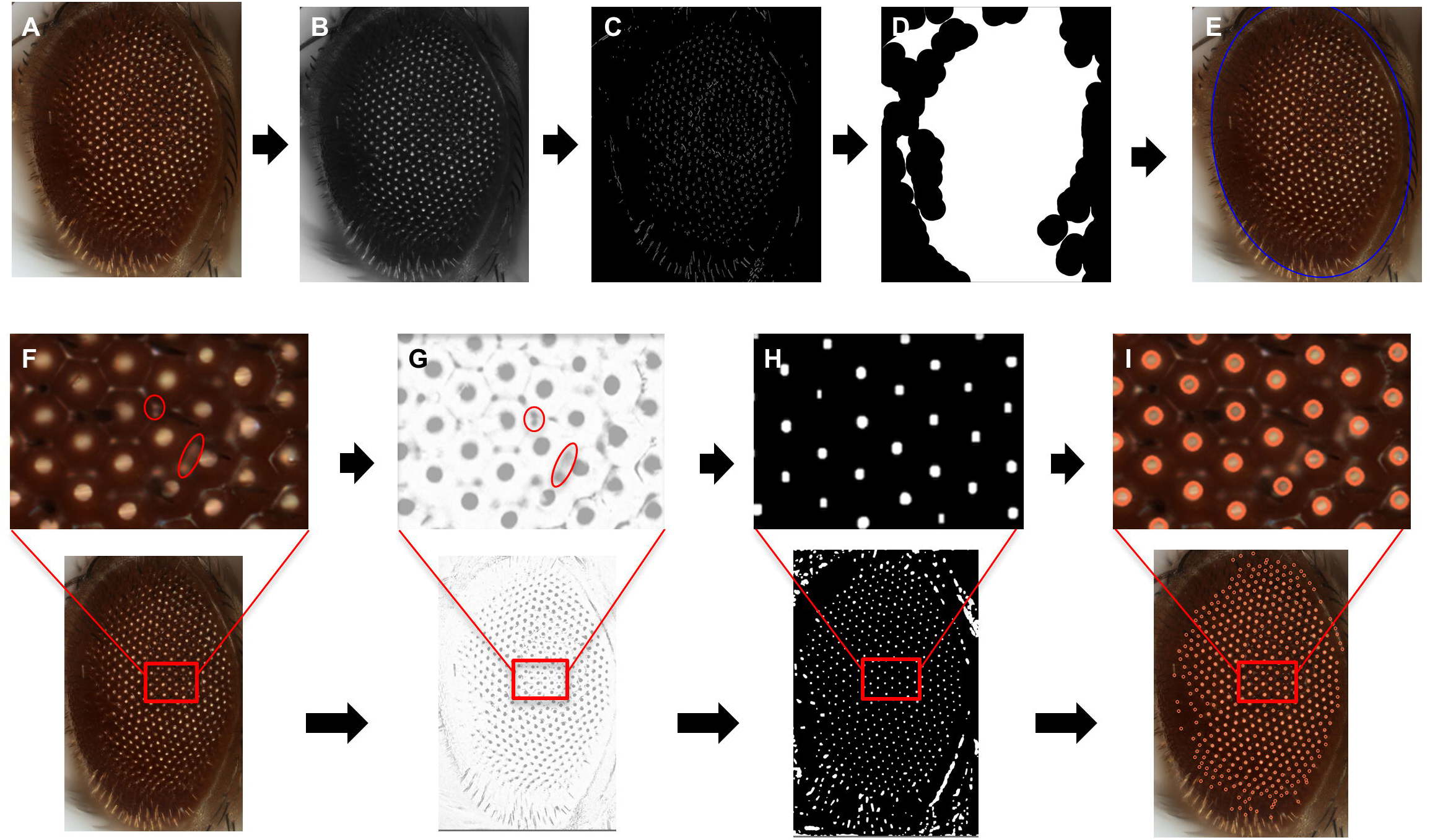
A computational strategy for automated assessment of *Drosophila melanogaster* eye morphology. Eye localization in a bright-field microscope image is carried out by first converting the original image (A) to grayscale (B), then morphological transformations are applied to suppress the background, followed by edge detection to identify an approximate region with ommatidial cluster (C), and finally morphological closing operation localizes the eye area (D), giving the final output image with the eye area localized (E). For detection of the ommatidial center, the original bright-field image (F) is converted to grayscale and inverted (G). Multiple filters and transformation operations further enhance the contrast of the inverted image and eliminate the noise from the ommatidial boundary (H). The centers of the bright spots due to light reflection are considered as ommatidial centers (I).

**Figure 2.**
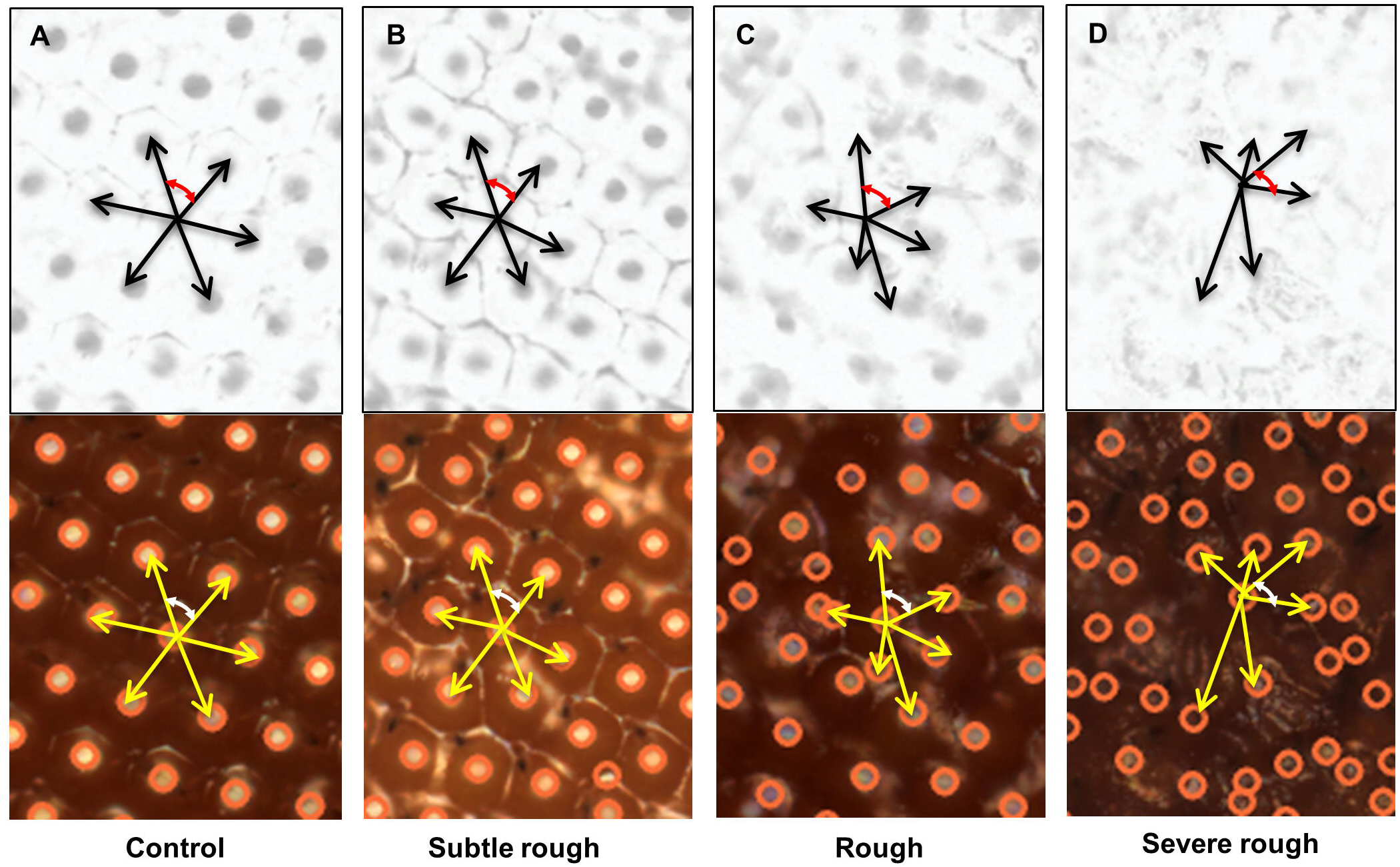
Detection of ommatidial centers and ommatidial disorderliness calculation in different classes of eye phenotypes. Grayscale inverted and bright field images of (A) Wild type control eye (B) subtle rough eye (C) rough eye and (D) severe rough eye phenotypes are shown. The ommatidial centers were accurately detected in all the four categories of eye phenotypes. The six local vectors are represented by black (for grayscale/inverted) and yellow arrows (for bright-field images). The red and white arrows represent the angle between two adjacent vectors in grayscale/inverted and bright-field images, respectively. Note the differences in the lengths of the local vectors and the angles between them in different classes of eye phenotypes.

### Assessing the performance of Flynotyper

We tested our algorithm on adult eye images from 21 lines with RNA-interference (RNAi) mediated eye-specific (GMR-GAL4/UAS-RNAi) knockdown of fly orthologs of human genes associated with neurodevelopmental disorders including *dpten, kismet, dube3a, prosap, arm, caps, para, rk, nrx-1, mcph1, tpc1, and eph* (**Table S1**). We hypothesized that these genes when knocked down in the eye would show varying levels of severity in eye phenotypes based on their level of knock down. These neurodevelopmental genes were chosen after curating the mutations observed in exome sequencing studies of autism (25), intellectual disability (26), schizophrenia (27, 28), and epilepsy (29), their roles in CNS development (30, 31), and their conserved functions in model organisms such as mouse and fly. Further the genes were shortlisted based on the presence of fly orthologs, high sequence identity, and availability of RNAi stocks.

A wide range of defects in eye morphology was observed for these 21 fly lines compared to a control line derived from the same genetic background but not expressing an RNAi construct (**Figure 3A-N**). Based on visual assessments of two independent reviewers, we manually ranked the eye phenotypes from 1 to 10 based on severity, with rank 1 assigned to wild type-like and rank 10 for the most severe phenotype (**Table S2**). These ranks were further classified into one of the following four broad qualitative categories including wild type-like, subtle rough, rough, and very rough (**Figure 3O**). A significant correlation (Pearson correlation coefficient, r=0.99, p<0.001) was observed when the manually determined ranks were compared to the corresponding phenotypic scores obtained from Flynotyper (**Figure 3O**). For example, knockdown of *dpten, kismet, dube3a, prosap, arm, nrx-1, mcph1* and *caps* resulted in significantly high phenotypic scores compared to control flies (student t test, corrected two-tailed p<0.001), indicative of a very rough eye phenotype (**Figure 3P, Table S3**). Similarly, lower phenotypic scores were observed for knockdown of *para, eph, tpc1* and *rk*, in agreement with the subtle rough eye phenotype observed in these flies (student *t* test, corrected two-tailed p<0.001). The phenotypic scores were also robust in distinguishing other qualitative categories of eye phenotypes such as glossy, necrotic and crinkled eyes (**Figure S3**). Notably, scores for the glossy eye phenotypes mapped well within these categories based on the extent of ommatidial fusion. For example, the glossy eyes with *dpten* knockdown was severe and scored similar to that of very rough eye, while glossy eyes with *kismet* knockdown scored similar to the rough eye. For all the fly lines tested, ten or more eye images were used to run on Flynotyper.

**Figure 3.**
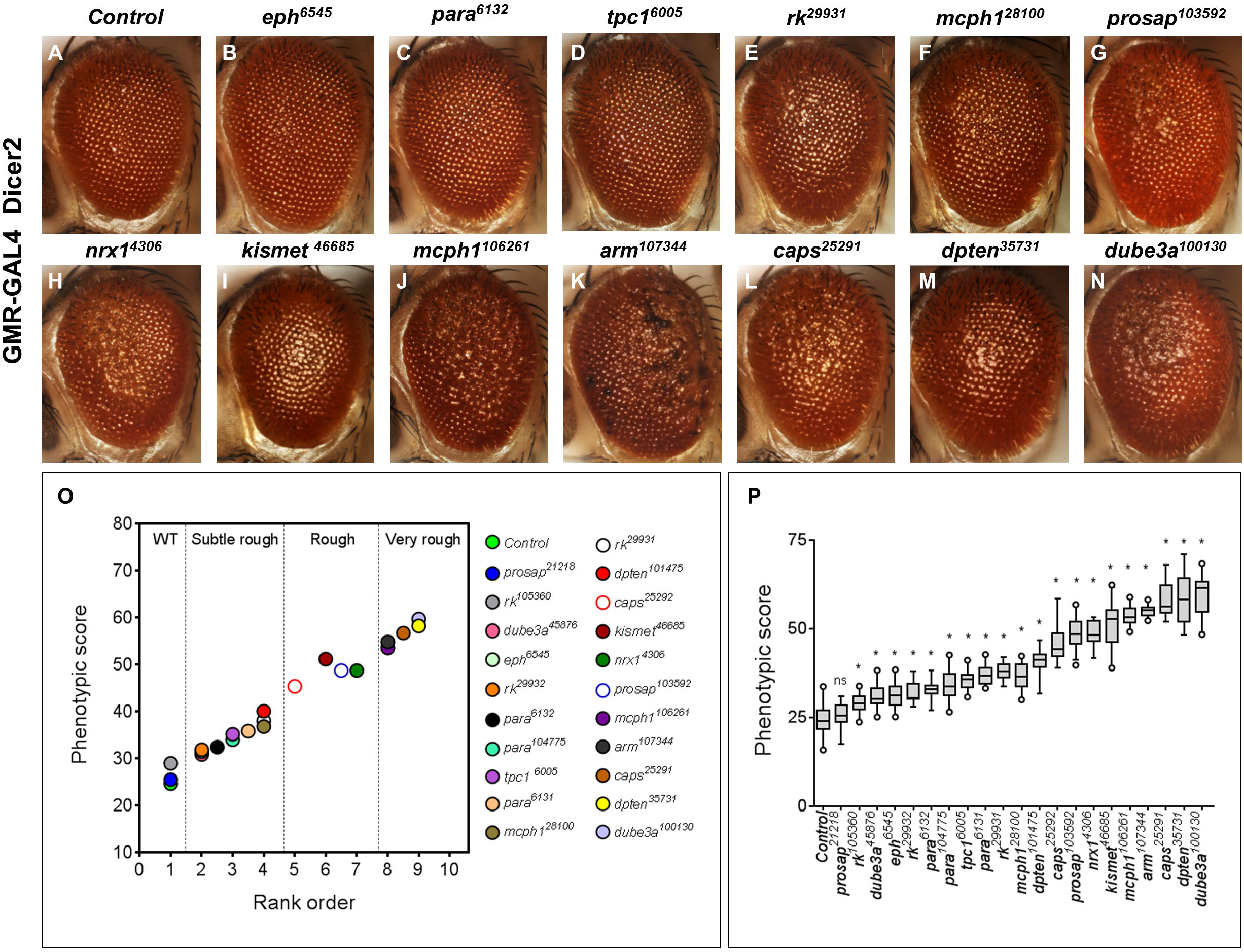
Analysis of *Drosophila* orthologs of human neurodevelopmental genes. (**A-N**) Representative bright-field microscope images of fly eyes displaying eye-specific knockdown of *tpc1, eph, para, rk, mcph1, prosap, nrx-1, kismet, arm, caps, dpten* and *dube3a* genes from flies reared at 30°C. Eyes of GMR-GAL4; Dicer2/+ control flies show normal ommatidial organization, while the eyes of flies with GMR-GAL4 driven RNAi knockdown of the 12 genes show disruption in the morphology of the eye. Note the variation in the severity of the eye phenotype for different genes. (**O**) Graph representing the mean phenotypic scores of control flies compared to the mean phenotypic scores for 21 RNAi lines with knockdown of neurodevelopmental genes (n=9 to 30). The rank order of these fly lines was significantly correlated with the phenotypic scores (Spearman correlation coefficient=0.99, p=1.2×10^−19^). (**P**) Graph representing phenotypic scores of the 21 RNAi lines with GMR-GAL4 at 30°C is shown. The number of images analyzed for each genotype ranged from 9 to 30 (median of 20.5). Comparisons were made between each of the gene knockdowns to controls using a student *t* test (*represents corrected two-tailed p<0.001).

Flynotyper phenotypic scores also allowed us to distinguish between genotypes within a specific phenotypic category. While all of the different *para* RNAi lines (*para^6131^, para^6132^*, and *para^104775^*) showed subtle rough eye phenotype, our algorithm was able to accurately identify differing degrees of severity imposed by each RNAi construct (**Figure S4**). Further, phenotypic scores were consistent with rank order even when different numbers (n=150, 200, 250, 300, 350) of ommatidia were used for the analysis (**Figure S5A**). In the Flynotyper program, we chose N=200 as the default number of ommatidia, since at this value the phenotypic distinction was robust with less variation in phenotypic scores between genotypes within and across ranks (**Figure S5B**).

We also evaluated the robustness of Flynotyper at different resolutions of images (600x800, 1200x1600 and 1600x2400) taken using a bright field microscope. Although greater number of ommatidia was detected for images acquired at higher resolutions (1200x1600 and 1600x2400) compared to those at a lower resolution (600x800), we found that the phenotypic scores were similar at all three resolutions tested (**Figures S6**). Thus our method can reliably quantify eye morphology at different counts of ommatidia and images taken at different resolutions. Traditionally, a SEM image is considered to be of higher quality than a bright field image, due to its ability to resolve finer details of the specimen. To determine the performance of our framework for SEM and bright field images, we compared the phenotypic scores for the same genotypes using images obtained from these two techniques (**Figure S7**). A positive correlation (Pearson correlation coefficient, r=0.95, p=0.011) was observed between the phenotypic scores obtained from SEM images to that from bright field images (**Figure S7C**). Thus Flynotyper processes images taken using SEM and bright field microscopy with similar accuracies.

We further performed a sensitivity analysis to determine the minimum number of eye images required for Flynotyper to accurately quantify and distinguish between the different categories of phenotypes. We chose one control and four genotypes, one from each category (control like, subtle rough, rough and very rough), and imaged ten eyes for each category and calculated their phenotypic scores. We then performed sampling with replacement with all possible combinations for n=3 to n=10 (where n is the number of eye images). Next we calculated the mean phenotypic score for each combination of eye images and generated plots for distribution of all possible mean scores. As expected, the spread of the distribution decreased as ‘n’ increased (**Figure S8A**). While no difference in the distribution of mean phenotypic scores was observed between controls and control-like eye images, significant differences (p<0.001, Mann-Whitney test) were observed when each category was compared with each other and with the control eye images for sample size as low as n=3 (**Figure S8B**).

### Testing dosage sensitivity of neurodevelopmental genes using Flynotyper

Next, we applied this method to test its ability to detect subtle differences in phenotypic effects caused by changes in the gene dosage of neurodevelopmental genes. We generated flies with different levels of mRNA for the same gene by either using different RNAi lines for the same gene or by rearing flies at different temperatures (28°C and 30°C). We performed qPCR experiments using fly heads for a subset of ten RNAi lines to confirm a reduction in the mRNA levels of the targeted genes (**Figure 4, Table S4**). The different RNAi lines of a gene were targeted to different sequences of the gene and may vary in their efficiency to knockdown, thereby resulting in different levels of mRNA. For example, knockdown of *pten* in two different RNAi lines, *pten^101475^* and *pten^35731^* resulted in mRNA expression of 55% and 25%, respectively. The phenotypic score of *pten^35731^* was higher than that from *pten^101475^*, in agreement with the level of knockdown. For the seven genes tested, the expression level was lower in flies reared at 30°C than in those reared at 28°C, due to the temperature-dependent effect of the UAS-GAL4 expression system (**Figure 4A, Table S5**). The phenotypic scores of flies reared at 30°C were correspondingly higher than those reared at 28°C, indicating a gene dosage-dependent increase in the severity of eye phenotypes for these genes (**Figure 4B**). In fact, Flynotyper was also able to identify subtle changes in the phenotype due to dosage alteration where visual assessment failed. For example, a significant difference in phenotypic scores was observed for *arm^107344^* flies reared at 28°C and 30°C and expressing 47% and 17% of mRNA, respectively (student *t* test, corrected two-tailed p=4.09×10^−18^).

**Figure 4.**
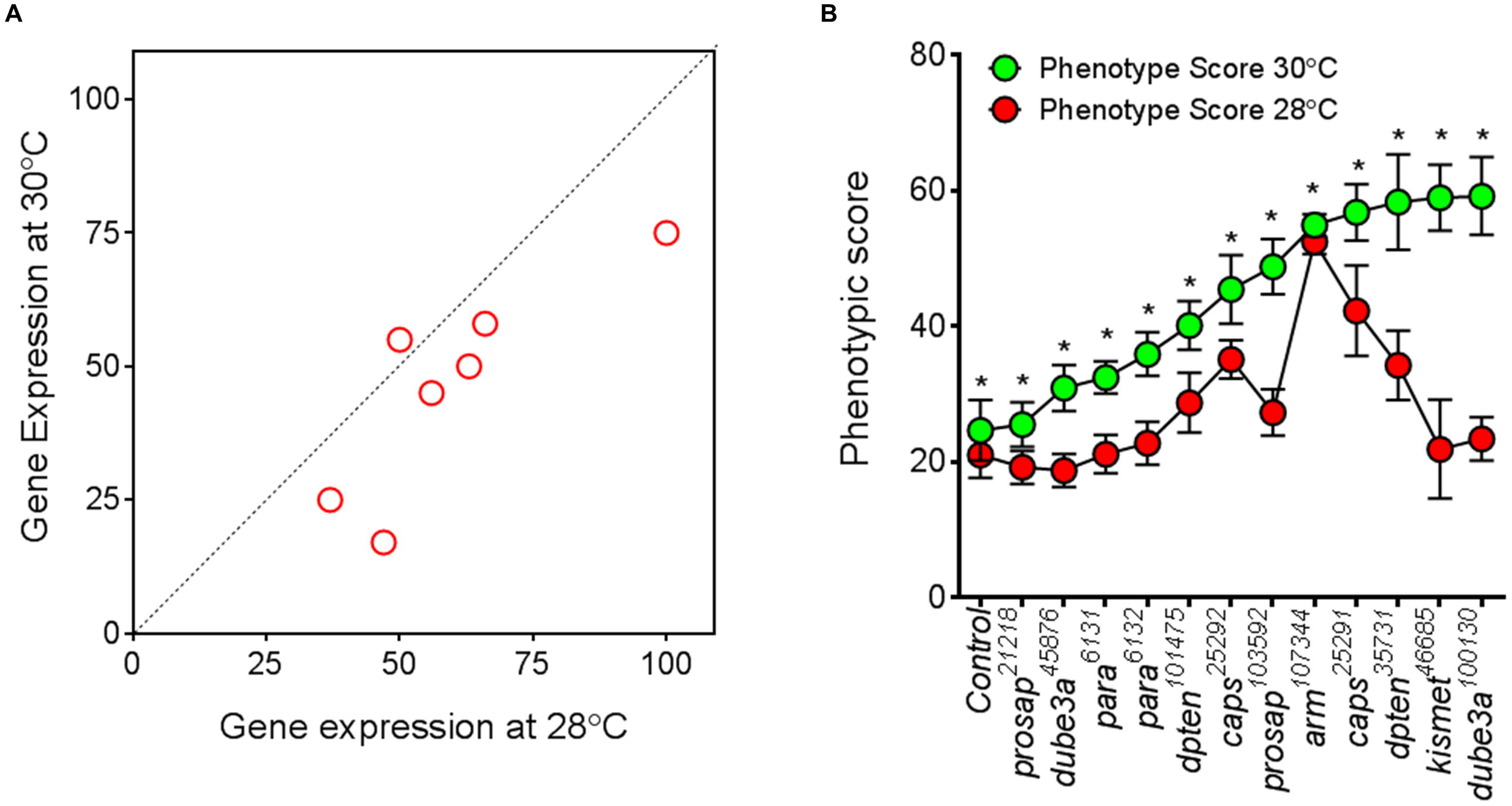
Quantitative assessment of eye phenotypes due to gene-dosage alteration. (**A**) Plot of mean values of gene expression of RNAi lines with eye specific knockdown at 28°C vs. 30°C. Note that each experiment was conducted in triplicates and then repeated using a fresh preparation of RNA and cDNA synthesis. Gene expression of flies reared at 30°C is lower than that at 28°C due to increased RNAi mediated knockdown at 30°C (note that most red circles are below the black diagonal line), indicating a temperature dependent effect of the UAS-Gal4 system. (**B**) A graph representing phenotypic scores for knockdown of *prosap, dube3a, para, pten, arm, caps* and *kismet* at 30°C and 28°C is shown. A dosage dependent increase in severity was observed for all the genes tested. Asterisks (*) show significant p-values (student *t* test, corrected two-tailed p<0.001) when phenotypic scores at 30°C was compared to that at 28°C. The number of images processed for each genotype ranged from n=14 to n=30. A complete list of statistical analysis and n numbers for each of the genotype assessed is presented in **Table S5**.

### Validation of Flynotyper using images from independent studies

To validate the robustness of our software for images generated using different image acquisition set ups (both bright field and SEM), we tested Flynotyper on adult eye images from six independent studies (both published and unpublished). We first tested the bright field images of eyes overexpressing *C9orf72* pure repeats and RNA only (RO) repeats (**Figure 5A & B**) (23). Expanded repeats in *C9orf72* is the most common genetic cause of frontotemporal dementia and amyotrophic lateral sclerosis (32-34). While expression of 36 and 103 pure repeats caused neurodegeneration in the fly eye, 36 RO repeats did not show any degeneration similar to 3 pure repeats, and 288RO and 108RO repeats showed mild degeneration, in agreement with the original assessment (23). We then tested a group of representative SEM images from a functional genetic screen for modifiers of tau-induced neurotoxicity (19). We were able to accurately identify and classify the different suppressors and enhancers of the human wild type full-length *tau* overexpression. While *cana, Klp61F* and *par1* enhanced the *tau* rough eye phenotype, *frc, ksr* and *sgg* suppressed the rough eye phenotype (**Figure 5C & D**). We also compared the phenotypic scores for each of these modifiers with the volumetric analysis reported in the published study (19), and observed a negative correlation (Pearson r=-0.71, twotailed p=0.049) with higher phenotypic scores corresponding to lower eye volumes (**Figure S9**). We also validated Flynotyper on images obtained from four additional studies, including a genetic screen for interactors of *dFoxO* (20) (**Figure S10A & B**), a screen for interactors of *DJ-1*, a gene mutated in familial Parkinson's disease (**Figure S10 C & D**), a positional scanning combinatorial peptide library screen to identify molecules that reduced CUG-induced toxicity in a *Drosophila* model of myotonic dystrophy type 1 (DM1) (21) (**Figure S11**), and a classical study of a genetic screen for interactors of *Egfr* using P-element insertions (22) (**Figure S12**). In the analysis of eye images from all these studies, Flynotyper was robust in accurately quantifying the severity of the eye phenotypes and concordant with the results from the original assessments.

**Figure 5.**
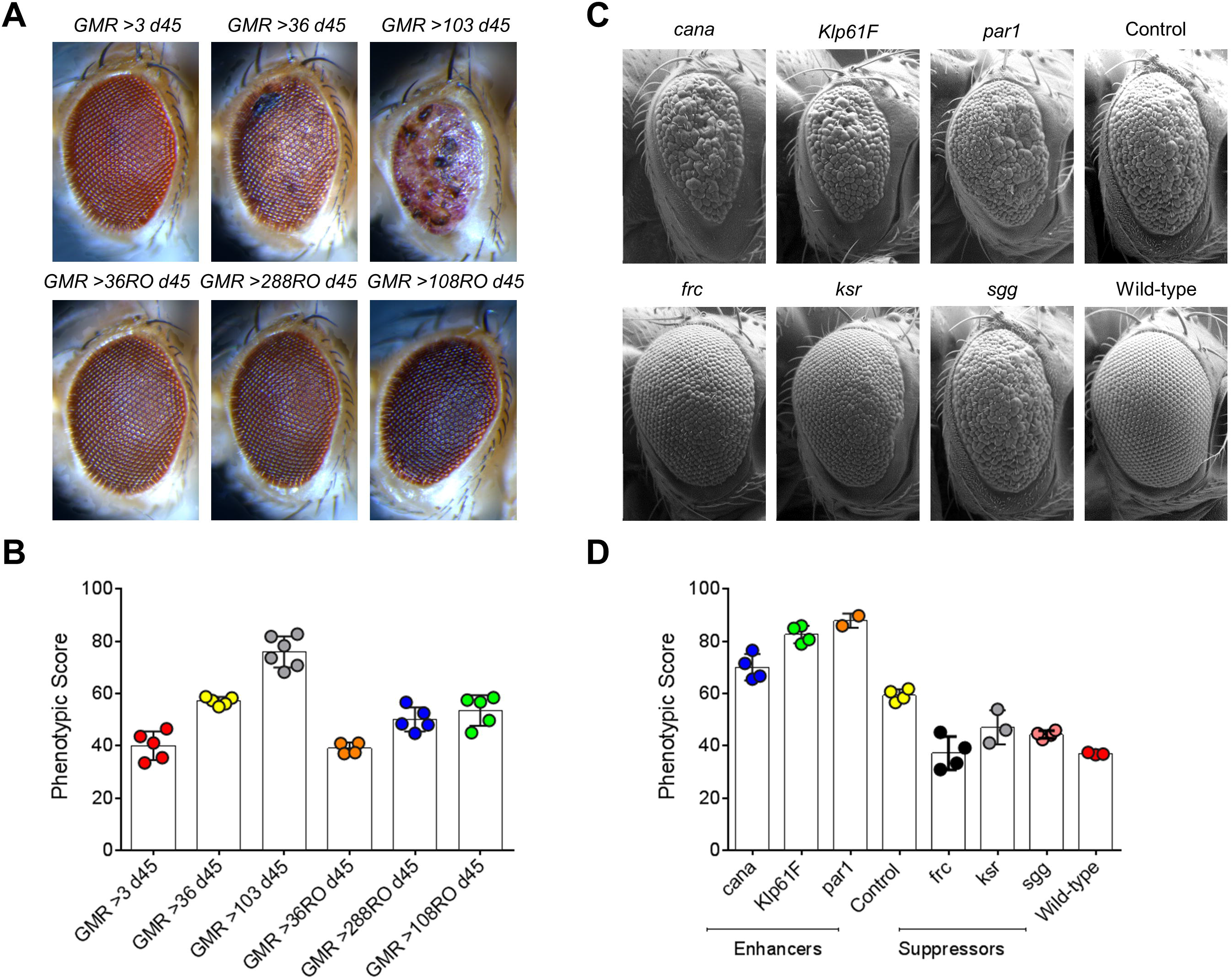
Analysis of eye images obtained from independent studies. **(A)** Bright-field microscopy images of representative *Drosophila* eyes overexpressing *C9orf72* pure or RO repeats using the GMR-GAL4 driver, imaged on day 45. While 3 pure repeats had no effect, 36 pure repeats were toxic and 103 pure repeats showed more overt toxicity. 36 RO repeats had no effect and 108 RO and 288 RO repeats showed mild effect. **(B)** A graph representing the phenotypic scores of *C9orf72* pure or RO repeats using the GMR-GAL4 driver is shown. The phenotypic scores are concordant with the visual assessment of the eye phenotypes (23). The number of images used for these assays were n=4 for GMR >36RO d45, n=5 each for GMR >3 d45, GMR >36 d45, GMR >288RO d45, and GMR >108RO d45, and n=6 for GMR >103 d45. **(C)** Scanning electron microscope images of genetic modifiers of tau-induced neurotoxicity. The control listed is *w^1118^/+;gl-tau/+*. All other panels, except wild type, contain one copy of *gl-tau* transgene *in trans* to one disrupted copy of the gene listed in the panel. **(D)** A graph representing the phenotypic scores of wild type, control, and the three enhancers and suppressors of *w^1118^/+;gl-tau/+*, is shown. The number of images used for these analyses were n=2 for par1, n=3 each for wild type and *ksr*, n=4 each for control, *cana, Klp61F, sgg*, and *frc*.

### Discovery of novel interactors of *sine oculis*

To illustrate the utility of our method in modifier screens, we used SEM images of candidate genetic interactors of *sine oculis (so)*, a *Drosophila* gene involved in eye development (35-37). Overexpression of *so* using ey-GAL4 and GMR-GAL4 results in a rough eye that is also significantly smaller in size compared to the wild type eye. Two genetic screens using 425 deficiency lines were performed to identify regions of the *Drosophila* genome that potentially interact with *sine oculis* (**Figure 6**). A total of 16 novel modifiers were identified by qualitative assessment, when flies overexpressing *so* with ey-GAL4 and GMR-GAL4 were each crossed to 425 deficiency lines (**Table S6**). These modifiers varied in their level of suppression or enhancement of the *so* overexpression eye phenotype, and could be broadly classified as mild, moderate, and strong modifiers. We first assessed the phenotypic scores of flies overexpressing *so*, using ey-GAL4, with or without the candidate modifiers and compared the effect of modifiers using student *t* test (**Table S7, Figure S13**). We identified four enhancers and four suppressors of *so* and used Flynotyper to accurately classify these candidate modifiers based on their effects. For example, among the tested deficiency lines, BL25005 was identified as a mild suppressor, BL8925 and BL2366 as moderate suppressors, and BL34665 as a strong suppressor (**Figure 6B-E, M**). Although the phenotypic scores of BL34665 and BL2366 are similar, BL34665 can be classified as a strong suppressor since the eye area of BL34665 is similar to that of wild type. We note that while our software does not directly calculate the size of the fly eye, a negative correlation exists between the size of the eye and the phenotypic score, as the calculation also accounts for the number of detectable ommatidia. Further, methods to quantify the eye size are also available, which can be incorporated into the framework of ImageJ along with Flynotyper (38-40). Similar phenotypic assessments led to classification of BL7659 as a mild enhancer, BL18322 as a moderate enhancer and BL27378 and BL7689 as strong enhancers (**Figure 6M**). Assessment of phenotypic scores of flies overexpressing so, using GMR-GAL4, with or without the candidate modifiers led to the identification of four additional suppressors and four enhancers of *so* (**Figure 6H-L, N, Table S7**). Among the tested deficiency lines, BL8674, BL7144, BL727 and BL3347 were identified as strong suppressors, BL3520 as a mild enhancer and BL1931, BL2414 and BL442 as strong enhancers (**Figure 6N**). Thus our method can be used to analyze the genetic interactions and also accurately classify the genetic modifiers strong and weak interactors.

**Figure 6.**
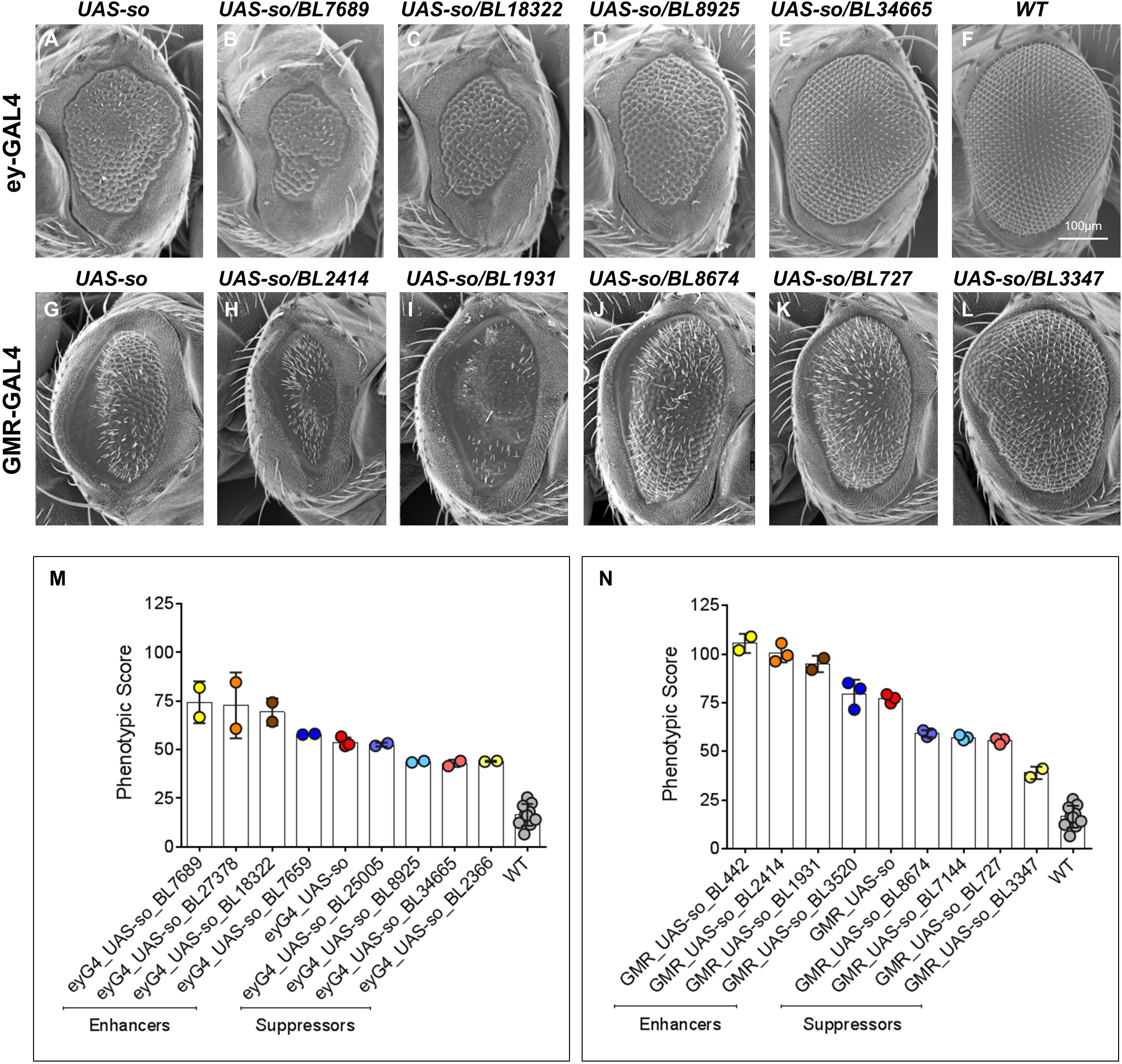
Genetic modifiers of *sine oculis*. (**A-L**) Scanning electron microscope images of adult compound eyes are shown. **(A)** ey-GAL4/UAS-*so*. Note that the eye is rough and smaller in size. **(B-C)** Enhancers of the rough eye phenotype due to overexpression of UAS-*so* with ey-GAL4 are shown. **(D-E)** Suppressors of the rough eye phenotype due to overexpression of UAS-*so* with ey-GAL4 are shown. Note that BL34665 rescues both the small eye and the rough eye phenotype of ey-GAL4/UAS-*so*. **(F)** Wild type (WT). **(G)** GMR-GAL4/UAS-*so*. **(H-I)** Enhancers of the rough eye phenotype due to overexpression of UAS-*so* with GMR-GAL4 are shown. **(J-L)** Suppressors of the rough eye phenotype due to overexpression of UAS-*so* with GMR-GAL4 are shown. (**M**) A graph representing the phenotypic scores of WT, ey-GAL4/UAS-*so*, and the four enhancers and four suppressors of ey-GAL4/ UAS-*so*, is shown. The number of images processed for each genotype ranged from n=2 to n=10. (N) A graph representing the phenotypic scores of WT, GMR-GAL4/UAS-**so** and the four enhancers and four suppressors GMR-GAL4/UAS-*so*, is shown. The number of images processed for each genotype ranged from n=2 to n=10. Note that the phenotypic scores clearly distinguish the effect of different suppressors and enhancers on the eye phenotype.

## Discussion

The *Drosophila melanogaster* eye has been used as a model to study various developmental processes (4, 8). Classical studies on *Drosophila* have established a line of research using the fly eye as an experimental system for studying genetic effects (5, 7, 41, 42). For decades, the fly eye has been used as a system for functional screening of genes and genetic interactions involved in basic cellular processes, neuronal development and degeneration, and common complex diseases such as cancer and diabetes. For example, Alzheimer's disease was successfully studied using the *Drosophila* eye by transgenic expression of human β-amyloid and β-secretase genes. These transgenic flies displayed age-dependent neurodegeneration and β- amyloid plaque formation, both of which were rescued by addition of inhibitors of β-secretase and γ-secretase (43). Similarly, *Drosophila* eye models have been used to characterize genetic interactions. For example, He and colleagues generated a fly model of protein-misfolding disease by misexpressing human proinsulin protein in the eye and identified novel genetic interactors (44). Another genetic screen for CLASP interactors using the *Drosophila* eye resulted in the identification of 36 genetic modifiers (45). However, most studies using the fly eye are qualitative in nature, involving visual inspection of the phenotypes, and manual scoring of genetic interactors. In order to address this issue of a lack of highly sensitive and quantitative assays for scoring eye phenotypes, we developed a computational method for quantitative assessment of eye morphology in fruit flies. We have implemented our algorithm as a software package called Flynotyper, which is available as an open source software.

The following features of Flynotyper are noteworthy (**Table S8**). First, our algorithm is highly sensitive and robust across different types of eye morphological defects, including rough eye, glossy eye, crinkled eye, and necrotic eye. The results of the performance of Flynotyper indicate that this method accurately detects even subtle alterations in eye morphology and provides a quantitative measure that can distinguish between varying severity of eye defects. This feature of the algorithm has allowed us to quantify the effect of dosage alteration of key neurodevelopmental genes. For example, Flynotyper accurately distinguished the severity of eye phenotypes for knockdown of *prosap, dube3a, para, pten, caps, arm* and *kismet*, at 28°C and 30°C. In fact the change in the eye phenotype due to dosage alteration of *arm* at 28°C and 30°C was indistinguishable by visual assessment, but Flynotyper was able to identify the subtle changes. The subtle alterations in the eye phenotypes between the different *para* and *pten* RNAi lines were also accurately detected by Flynotyper. Second, we have validated the robustness of Flynotyper using different images generated using different image acquisition set ups. We tested Flynotyper on adult eye images from six independent studies (both published and unpublished) and found that our analysis was concordant with the original assessments. In fact we found a negative correlation of phenotypic scores with the volumetric assessments reported by Ambegaokar and Jackson (19), with higher phenotypic scores corresponding to lower eye volumes. Third, Flynotyper is capable of quantifying eye images taken from both bright field and SEM and is equally efficient in analyzing images of different resolutions. Performing two genome-wide screens using deficiency lines, we obtained a total of 16 novel candidate modifiers of an eye development gene, *sine oculis*. Flynotyper was able to accurately quantify the effect of its enhancers and suppressors of *so* and also classify them as mild, moderate or strong modifiers. Finally, this tool is fast (3 seconds to process a 1800×2400 resolution image) and user friendly, rendering it ideal for use by non-experts to quickly quantify images from multiple genotypes associated with various developmental and degenerative processes.

While we provide the first quantitative method for assessment of eye morphology and demonstrate its robustness using images from independent studies, in its current version, Flynotyper cannot automatically quantify certain eye phenotypes such as bristle integrity, loss of pigmentation, overall eye size, ommatidial size, and altered photoreceptor integrity that is not visible on the surface (**Table S8**). We note that Flynotyper is not an alternative to visual inspection or other phenotypic methods of the eye. Rather, it is a tool that will complement other existing assays, such as the thin sectioning of the eye (46), pseudopupil analysis (47) and semiautomated methods to measure eye size (38-40), neurodegeneration (48, 49), and pupal eye patterning (50), to enable more accurate measurement of genetic effects (**Figure S14**). The strengths of Flynotyper will enhance the sensitivity of studies that use rough eye phenotypes for understanding the effects and interactions of genes involved in various biological processes.

*Drosophila* research has laid the ground for several highly impactful discoveries in neuroscience and continues to do so (3). With new technologies facilitating more accurate genetic manipulation (51-54) and sequencing studies providing an unprecedented number of candidate genes, there is now an increasing need for functional genomics. Our study will have a broader impact on quantitative genetic screens and emphasizes the use of fly models for modeling human diseases. Our tool is an open source software available for free download at http://flynotyper.sourceforge.net.

## Materials and Methods

### *Drosophila* stocks

The conditional knockdown of specific genes was achieved with the UAS-GAL4 system (55), using w;GMR-GAL4; UAS-Dicer2 (Zhi-Chun Lai, Penn State University) and the UAS-RNAi transgenic lines. The following RNAi fly stocks from Vienna Drosophila Resource Center (Austria, Vienna) (56) were used in this study: UAS-*ube3a^RNAi^* (VDRC# 100130, 45876), UAS-*prosap^RNAi^* (VDRC# 103592, 21218), UAS-*caps^RNAi^* (VDRC# 25291, 25292), UAS-*kismet^RNAi^* (VDRC# 46685), UAS-*para^RNAi^* (VDRC# 6132), UAS-*pten^RNAi^* (VDRC# 101475, 35731), UAS-*arm^RNAi^* (VDRC# 107344), UAS-*rk^RNAi^* (VDRC# 105360, 29931, 29932), UAS-*tpc1^RNAi^* (VDRC# 6005), UAS-*eph^RNAi^* (VDRC# 6545), UAS-*nrx1^RNAi^* (VDRC# 4306) and UAS-*mcph1^RNAi^* (VDRC# 106261, 28100). The following stocks were used in studying genetic interactions, ey-GAL4 and GMR-GAL4 with UAS-so and the Bloomington Stock Center Drosophila deficiency kit (Bloomington Stock Center) (54). All stocks and crosses were cultured on conventional cornmeal/sucrose/dextrose/yeast medium at 25°C unless otherwise indicated. A list of all genotypes obtained from the deficiency screen is given in **Table S6**.

### Quantitative real-time PCR

We assessed mRNA expression using quantitative real-time PCR (qRT-PCR) by isolating RNA from fly heads expressing RNAi knockdown of specific genes. Groups of 40-50 female flies were frozen in liquid nitrogen and stored at -80°C. For RNA extraction their heads were separated from bodies by repetitive cycles of freezing in liquid nitrogen and vortex mixing. Total RNA was isolated using TRIZOL (Invitrogen) and reverse transcribed using qScript cDNA synthesis kit (Quanta Biosciences). Quantitative RT-PCR was performed using an Applied Biosystems Fast 7500 system with SYBR Green PCR master mix (Quanta Biosciences). All SYBR green assays were performed in triplicate and normalized to *rp49* mRNA expression. Each qRT-PCR experiment was repeated twice with two independent RNA isolations and cDNA syntheses. A list of primers used for these experiments is provided in **Table S4**.

### Eye imaging using bright field microscope

For light microscope imaging of adult eyes, the 2-3 day old flies (GMR;Dicer2 >UAS-RNAi), reared at 28°C and 30°C, were immobilized by freezing at -80°C and then mounted on blu-tack (Bostik Inc, Wauwatosa, WI). These flies were then imaged using Olympus BX53 compound microscope with a LMPlanFL N 10X 0.25 NA air objective (Olympus, Tokyo, Japan), at 0.5X magnification and a z-step size of 12.1μm. Images were captured with CellSens Dimesion software (Olympus Optical) and the slices stacked using Zerene Stacker (Zerene Systems, USA). All images shown in figures are maximum projections of consecutive optical z-sections.

### Eye imaging using Scanning Electron Microscope (SEM)

Flies aged 2-3 days old (GMR; Dicer2 >UAS-RNAi reared at 30°C) were anesthetized with CO_2_ and dissected midway through the abdomen and fixed overnight at 4°C in 0.2M Sodium cacodylate buffer containing 2.5% glutaraldehyde. The preparations were then washed three times with 0.1M Sodium cacodylate buffer for five minutes each at room temperature. The samples were then washed through ethanol series (50%, 70%, 85%, 95%, and 100%) and critically point dried using Leica CPD300 at the Penn State Microscopy and Cytometry Facility (The Pennsylvania State University, University Park, PA). The dried flies were mounted onto carbon taped SEM stubs, and sputter-coated with a 25-nm-thick gold coat. Samples were then imaged using Zeiss Field Emission-SEM (Carl Zeiss AG, Oberkochen, Germany).

### The Flynotyper computational method

**Detection of *Drosophila* eye from ommatidial clustering:** We obtained high quality images of eyes from individual fly genotypes using light microcopy or monotonic images from SEM. The fly eye is not flat but convex in shape, and each ommatidium is also convex (**Figure 1A-E**). Therefore, in a bright field image, when light falls on each ommatidium, there is a reflection spot in its center. The whole eye area is then presented as a region enriched for such bright spots, which also leads to a high contrast between the eye and the background in the image. Conversely in a SEM image, there is a dark spot in the center of each ommatidium compared to the boundary, which might be due to variations in collection efficiency of the secondary electrons, owing to the convex shape of the ommatidium. First, the original images are converted into gray scales. A series of morphological transformations are applied to suppress the background and identify the fly eye based on the clustering of the ommatidia (24). The result of different morphological transformation depends on the different size of the matrix and what operation was applied. The small pixel matrix is then referred to as the structuring element (SE). As the quality of the final output extracted from the image depends on the size of the structuring element, we tested different combination of sizes of structural elements to optimize the pixel intensity of the circular shaped ommatidia (**Figure S15**). We performed a top-hat transformation to extract light objects from a dark background (57). As a result the background is suppressed as pixel values are subtracted from the input frame, reducing its intensity and enhancing the eye region in the process. Edge detection was then applied over the resultant image to identify an approximate region with a clustering of ommatidia. Following edge detection, we used a morphological closing operation to expand the boundaries of the foreground area in the image by filling in the surrounding background to connect clusters of all edge-detected regions. By detecting the largest connected area, we were able to localize the position of the *Drosophila* eye as a region with ommatidial clustering.

The images obtained from scanning electron microscope were processed differently as a low contrast between the foreground (eye) and the background limited the application of top hat transformation to suppress the background (57). In addition, the SEM images capture the details in the area surrounding the eye, thereby significantly increasing the background noise and thus hindering the accurate detection of the ommatidial cluster. To circumvent this issue, we applied a thresholding operation for SEM images (**Figure S16**). Thresholding operation is a basic segmentation method performed on a gray-scale image to separate out objects of interest from the image background (24). This separation is based on the difference in pixel intensity between the object and the background. By using the mean intensity of the image as a determined cutoff, we could separate the region of interest from the background. The subsequent steps of the method are the same for images taken using both light microscope and SEM.

**Detection of ommatidia by applying morphological transformation and searching local maxima:** In a digital image, ommatidia appear as roughly circular shapes also referred to as blobs. Due to the fusion of ommatidia observed in different genotypes, the boundary of each ommatidium can show non-uniform intensities. Our next step was to isolate each ommatidium as a single blob and remove the noise caused by the boundary (**Figure 1F-I**). The following steps were performed to extract ommatidia. We first enhanced the contrast between the ommatidia and the background. In a bright-field image, the contrast is between the reflection spot at the center of the ommatidium and the rest of the ommatidium, while in a SEM image, this contrast is between a darker center and lighter boundary of the ommatidium. Then, by applying top hat transformation on the enhanced image, the ommatidia were highlighted from the background (57). However, noises such as light reflection on the boundary between neighboring ommatidia also gets extracted. These noises were then removed by using the median filter. The median filter is used to blend the intensity of the pixels within a specified radius by calculating the median value of neighboring pixels in a specific structuring element and assigning that value to each pixel in the element. Thus, the background noise was removed by smoothing the pixels in the neighboring region resulting in bright blobs representing ommatidia. Because of the non-uniformity and reduced intensity of each blob, we further enhanced the contrast of the image using a combination of dilation and erosion operations. As a result, each ommatidium was isolated as a single blob and its center was localized by searching for local maxima within the fly eye. Using the centers of ommatidia, phenotypic scores were then computed based on the distance and the angle between the ommatidia. Our algorithm robustly detected the ommatidial centers in a variety of eye phenotypes qualitatively classified as wild type-like, subtle rough, rough and severe rough (**Figure 2**).

**Calculation of phenotypic score:** The *Drosophila* compound eye consists of a hexagonal array of packed ommatidia. In wild type compound eyes, ommatidia are ordered as mirror images that are radially symmetrical about an axis. The symmetric arrangement is disrupted in a defective eye resulting in an irregular hexagonal arrangement. Based on this principle, we determine groups of six local vectors, v, with direction pointing from each ommatidium to six surrounding ommatidia. We quantified the disorganization of the ommatidia by applying the principle of entropy, a measure of disorderliness, in which a perfectly radial symmetry resulted in zero entropy. To quantify the disorderliness of the fly ommatidia, we calculated phenotypic scores based on three measures including distance ommatidial disorderliness index (ODI_D_), angle ommatidial disorderliness index (ODI_A_), and ommatidial fusion index (Z). These indices essentially measure the level of disruption of symmetry of the ommatidia. The mathematical formulation to calculate ODI_D_ and ODI_A_ is described as follows:

As illustrated in **Figure 2**, distance ommatidial disorderliness index is defined as the difference between the lengths of each of the five local vectors, v_i_ (i=1…5), from the smallest vector, v_min_. Distance ommatidial disorderliness index for each ommatidium is calculated as

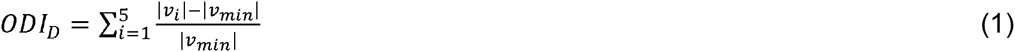

Similarly, angle ommatidial disorderliness index for each ommatidium is the difference among angles formed by pairs of adjacent local vectors to the smallest angle between the vectors. Thus, angle ommatidial disorderliness index for each ommatidium is calculated as:

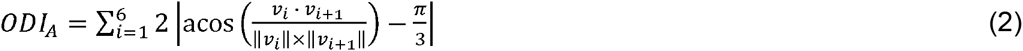

Here, v_i+1_ indicates the vector adjacent to v_i_ (Note the vector adjacent to v_6_ is v_1_).

The total ommatidial disorderliness index, ODI_T_, is the sum of distance and angle ommatidial disorderliness indices using the number of most ordered ommatidia (N, a tunable parameter) and is determined by equations (1) and (2) as given below:

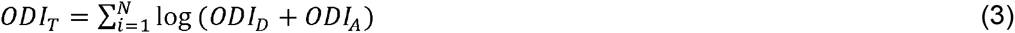

In this equation, N is the parameter defined by the user to be the number of most ordered ommatidia. If we let Fusion index Z represents the number of ommatidia being identified by the detection algorithm, then phenotypic score *P* is computed as followed using (4)

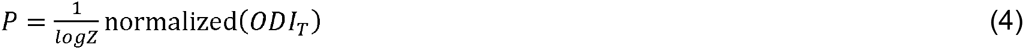

The ommatidia are rank-ordered based on their total ommatidial disorderliness index values and the distance to the center of the eye. As the phenotypic score is predominantly dependent on the ommatidial disorderliness index, higher phenotypic score represents increased disorderliness of the ommatidia and thus increased severity of the eye phenotype.

## Acknowledgements

The authors thank Drs. Michael Grotewiel, Shashikant Cooduvalli, Richard Ordway, Fumiko Kawasaki, Francesca Chiaromonte, Claire Reynolds, Scott Selleck, and Zhi-Chun Lai for useful discussions and comments on the manuscript. We thank the Microscopy and Cytometry Facility at the Huck Institute of Life Sciences, Penn State University for assistance with SEM images. This work was supported in part by a Basil O’Connor Award from the March of Dimes Foundation (#5-FY14-66) and a NARSAD Young Investigator Grant from the Brain and Behavior Research Foundation. The authors declare that no conflict of interest exists in relation to this work.

